# Effect of different forage-to-concentrate ratios on the structure of rumen bacteria and its relationship with nutrition levels and real-time methane production in sheep

**DOI:** 10.1101/585208

**Authors:** Runhang Li, Zhanwei Teng, Chaoli Lang, Haizhu Zhou, Weiguang Zhong, Zhibin Ban, Xiaogang Yan, Huaming Yang, Mohammed Hamdy Farouk, Yujie Lou

**Affiliations:** College of Animal Science and Technology, Jilin Agricultural University, Changchun, PR China; Jilin Academy of Agricultural Sciences, Changchun, PR China; Animal Production Department, Faculty of Agriculture, Al-Azhar University, Nasr City, Cairo, Egypt

**Keywords:** greenhouse gas, high-throughput sequencing, ruminant, rumen bacteria, rumen fermentation

## Abstract

Emission from ruminants has become the largest source of anthropogenic emission of methane in China. The structure of the rumen flora has a significant effect on methane production. To establish a more accurate prediction model for methane production, the rumen flora should be one of the most important parameters. The objective of the present study was to investigate the relationship among changes in rumen flora, nutrient levels, and methane production in sheep fed with the diets of different forage-to-concentration ratios, as well as to screen for significantly different dominant genera. Nine rumen-cannulated hybrid sheep were separated into three groups and fed three diets with forage-to-concentration ratios of 50:50, 70:30, and 90:10. Three proportions of the diets were fed according to a 3 × 3 incomplete Latin square, design during three periods of 15 d each. The ruminal fluid was collected for real-time qPCR, high-throughput sequencing and *in vitro* rumen fermentation in a new real-time fermentation system wit. Twenty-two genera were screened, the abundance of which varied linearly with forage-to-concentration ratios and methane production. In addition, during the 12-hour *in vitro* fermentation, the appearance of peak concentration was delayed by 26-27 min with the different structure of rumen bacteria. The fiber-degrading bacteria were positively correlated with this phenomenon, but starch-degrading and protein-degrading bacteria were negative correlated. These results would facilitate macro-control of rumen microorganisms and better management of diets for improved nutrition in ruminants. In addition, our findings would help in screening bacterial genera that are highly correlated with methane production.

## Introduction

Of the total methane emission in China, the emission from ruminants was estimated to be approximately 17%, turning them into the largest anthropogenic source of methane emission [1]. The emission of methane associated with agriculture is expected to see a significant increase. Therefore, new strategies were needed for reducing the emission and improving livestock productivity, which had been extensively studied and reviewed [2].

Rumen is the main site of methane production [3], which provided a habitat for a variety of microbes, including numerous species of bacteria, archaea, viruses, protozoa and fungi [4]. In the anaerobic environment of the rumen, several organic compounds present could eventually be decomposed into methane by a number of microorganisms [5]. The composition of ruminal microbiome was affected by different factors, such as age, breed, general well-being of the animal, its location as well as administration of feed and antibiotics [6–8]. Furthermore, feedstuffs were the main factors regulating the composition and functional patterns of ruminal microbiome [9–11]. Among the nutritional indices of diets, protein and energy levels were the major factors affecting the fermentation of ruminal microbiome [12]. Fibers, including hemicellulose and cellulose, were the main source of energy [13], which could be degraded into methane by the microbes present in the rumen. Leng and Nolan [14] pointed out that 80% of the nitrogen available to ruminal bacteria came from ammonia and 20% was derived from amino acids or oligopeptides. Therefore, the low content of ammonia promotes microorganisms to degrade other nitrogen sources in a diet with high forage-to-concentration ratios (F:C), which delays fermentation. Grovum and Leek [15] found that non-structural carbohydrates were degraded much faster than structural ones. Easily degradable carbohydrates provide energy and carbon sources for faster microbial fermentation and increase fermentation rate. Methane production can be affected by the above-mentioned factors.

In the current models established for the same rumen microflora to predict methane production, nutritional indicators had been used as parameters [16–21]. A large number of calibration parameters are required for the models to adapt to plentiful situations, thereby limiting the scope of these models.

Some mechanistic models considering the role of rumen microbes [22–23] had been established by the extrapolation of mathematical formula used by computers. Because of high operation cost, it is difficult to apply these models to actual production systems. Therefore, the application scope of these models will be greatly expanded if some important microorganisms can be related with the models using nutrient indicators as parameters.

Previous studies have indicated that archaea are the main microorganisms producing methane in the rumen [24]. However, other recent studies involving high-throughput sequencing have shown that change in methane production is irrelevant to archaea flora, but highly correlated with bacterial flora [25]. The main function of bacteria is to break down the nutrients in the feed into simple compounds and additional products used by animals, including hydrogen, carbon dioxide and volatile fatty acids which are raw materials for methane synthesis [3]. To establish better models for methane prediction with wide range of application, characteristic microorganisms should be screened from rumen bacterial communities to serve as effective parameters.

Simple devices for *in vitro* fermentation have been used to establish the prediction models [21]. However, in such cases, methane production could only be detected either at specific time points or at the final time point, and therefore, did not reveal the overall fermentation status well. Sun et al. [26] used a new real-time *in vitro* fermentation system to determine the methane production time course when they studied the effect of cysteamine hydrochloride and nitrate supplementation on methane production and productivity in cattle. This system makes it possible to determine a more subtle fermentation state. Thus, in exploring the relationship between rumen microbial structure and methane production, this system may provide more detailed reference data.

We hypothesized that the real-time methane production of sheep would be highly correlated with the abundance of bacteria in the rumen. The objective of this study was to investigate the relationship among different structures of bacterial flora in the rumen, dietary levels, and methane production, using the *in vitro* fermentation system. The genera of bacteria that showed high correlation with methane production were screened in order to serve as the reference for accurate prediction of methane production.

## Material and methods

### Ethics statement

All research involving animals was conducted according to Guide for the Care and Use of Laboratory Animals which was approved by the ethics committee of Jilin Agricultural University, P. R. China. The ethics committee of Jilin Agricultural University, P. R. China approved this study, and the approved permit number for this study is “JLAC20171104”.

### Animals and diets

A total of nine rumen-cannulated (cannulated at one year of age) hybrid sheep (Chinese merino fine wool sheep × Dorper sheep) were selected, which was 2 years old and whose average weight was 87.83 ± 8.11 kg. Randomly assigned to three groups, these sheep were separately fed at random. Jilin Agricultural University, Changchun, China prepared Guide for the Care and Use of Laboratory Animals which provided guidance for all animal-related procedures.

Chosen as the forage, *Leymus chinensis* was mixed with the concentrate in three proportions including 50:50 (L), 70:30 (M) and 90:10 (H). The composition and nutrition levels of the three diets based on the NRC [27] are shown in Table 1. The three diets were fed according to the 3 × 3 incomplete Latin square design over 45 d in three periods of 15 d each, including 14 d of pre-feeding and the 15th day for sampling. Three distinct flora structures were established under different treatments.

**Table 1.** Ingredients and nutrient compositions of diets

| Item | Treatments <sup>a</sup> |  |  |
| --- | --- | --- | --- |
|  | L | M | H |
| Ingredient (Fresh matter, g/kg) |  |  |  |
| *Leymus chinensis* | 500.0 | 700.0 | 900.0 |
| Corn | 237.5 | 137.5 | 37.5 |
| wheat bran | 118.7 | 68.7 | 18.7 |
| Soybean meal | 95.0 | 55.0 | 15.0 |
| Cottonseed meal | 23.8 | 13.8 | 3.8 |
| Calcium carbonate | 4.0 | 4.0 | 4.0 |
| Calcium hydrogen phosphate | 5.0 | 5.0 | 5.0 |
| Sodium chloride | 6.0 | 6.0 | 6.0 |
| Premix <sup>b</sup> | 10.0 | 10.0 | 10.0 |
| Composition <sup>c</sup> |  |  |  |
| DM (g/kg) | 887.4 | 889.2 | 891.1 |
| 105 °C DM (g/kg) |  |  |  |
| CP | 138.8 | 113.9 | 89.0 |
| EE | 25.7 | 29.7 | 33.6 |
| Ash | 32.8 | 34.6 | 36.3 |
| Starch | 267.1 | 185.8 | 104.6 |
| NDF | 296.2 | 398.6 | 501.0 |
| ADF | 149.6 | 221.4 | 293.1 |
| ADL | 31.8 | 43.1 | 54.5 |

DM, dry matter; CP, crude protein; EE, ether extract; NDF, neutral detergent fibre; ADF, acid detergent fiber; ADL, acid detergent lignin.
<sup>a</sup> L = forage-to-concentrate ratio 50:50; M = forage-to-concentrate ratio 70:30; H = forage-to-concentrate ratio 90:10.
<sup>b</sup> Provided per kilogram of premix: 80,000-145,000 mg of vitamin A, 20,000-39,000 mg of vitamin D, $\geq 700$ IU of vitamin E, 180-345 mg of Cu, 190-330 mg of Fe, 950-1,800 mg of Zn, and 350-650 mg of Mn.
<sup>c</sup> Calculated from the analyzed value of the dietary ingredients.

### Sampling and DNA extraction

Ruminal fluid (400 mL) was collected by pump and pre-warmed thermos before feeding (07:00 h) and saturated with CO_2_. Filtrated through a double-layered gauze, the collected fluid was used to measure the pH value to confirm the health of rumen. The fluid with the pH value between 5.5 and 7.5 from three sheep of one group was mixed. All the samples with 10 mL were respectively stored in sterile centrifuge tubes (without any treatment) with 2 mL at −80 °C for high-throughput sequencing. A total of 18 samples for the three diets were collected to have six replicates for each diet. Another 300 mL of ruminal fluid from each group was warmed to 39 °C for *in vitro* rumen fermentation right after sampling.

Microbial genomic DNA was extracted from all ruminal fluid samples with 220 mg using the methods of Murray and Thompson [28] and Zhou et al. [29]. Agarose gel electrophoresis was applied to confirm the successful extraction of DNA [30]. The qualified DNA continued to be tested for real-time quantitative polymerase chain reaction (qPCR) and high-throughput sequencing. A total of 9 samples for the three diets were collected and each sample was tested twice in order to have six replicates for each diet.

### Real-time qPCR for total bacteria, methanogens, protozoa and anaerobic fungi

Real-time qPCR were tested on Applied Biosystems StepOne™ Real-time qPCR System based on the methods of Denman and McSweeney [31]. The designed primers were shown in the Table 2.

**Table 2.**
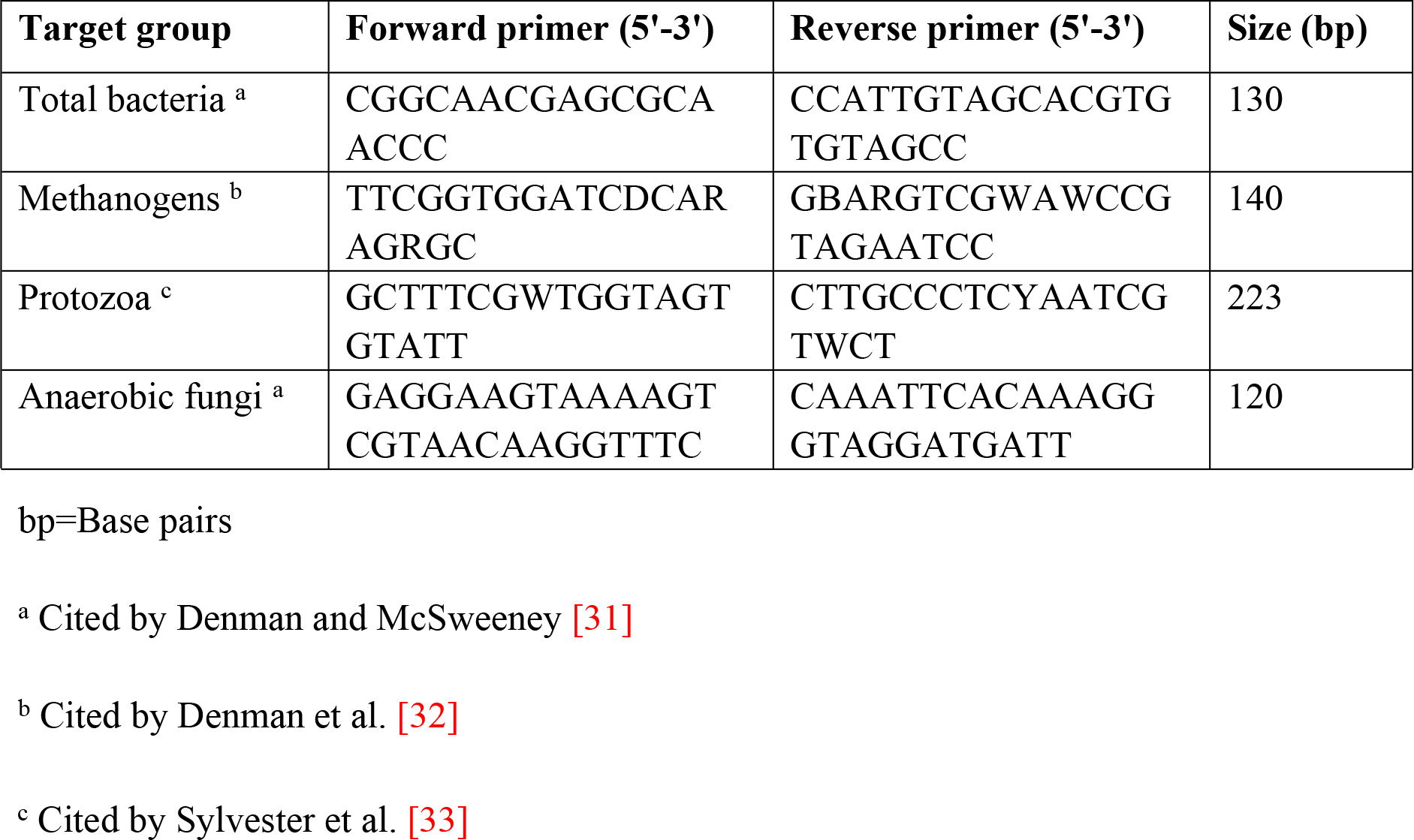
The primers for real-time qPCR assay

### PCR amplification of 16S rDNA, amplicon sequence and processing of sequence data

Based on previous comparisons [34–36], 16S rDNA had V_4_ hyper variable regions which performed PCR amplification for microbial genomic DNA extracted from ruminal fluid samples and were adopted in the rest of the study. PCR primers which flanked bacterial 16S rDNA’s V_4_ hypervariable region were designed. The forward primer with a barcode was 338F 5’-ACTCCTACGGGAGGCAGCAG-3’ while the reverse primer referred to 806R 5’-GGACTACHVGGGTWTCTAAT-3’ based on the approach of Fan et al. [37]. Below is the PCR reaction system (TransGen AP221-02, 20μL): 5×FastPfu Buffer, 2.5 mM dNTPs, Forward Primer (5μM), Reverse Primer (5μM), FastPfu Polymerase, BSA and Template DNA and ddH_2_O with 4μL, 2μL, 0.8μL, 0.8μL, 0.4μL, 0.2μL and 10 ng respectively were added up to 20μL totally. Below are PCR conditions: One pre-denaturation cycle, 27 denaturation cycles, annealing, elongation and one post-elongation cycle at 95 °C, 95 °C, 55 °C, 72 °C and 72 °C for 3 min, 30s, 30s, 45s and 10 min respectively. Separated on 1% garose gels, products of PCR amplicon were obtained by the extraction of gels. Sequencing only adopted PCR products which were void of contaminant bands and primer dimers by means of synthesis. Illumina MiSeq PE300 proposed the paired-end approach which was taken for the sequencing of barcoded V_4_ amplicons. The filtering of effective reads was based on the methods of [38–41]. Sequences with lower mean phred score (20 bp), equivocal bases and primer mismatching or sequence lengths below 50 bp were removed. The assembly of only the sequences which had an overlap above 10 bp and no mismatch was completed in accordance with their overlap sequence. Reads which were unable to be assembled were abandoned. Barcoded and sequencing primers were removed from the sequence which was assembled.

### Taxonomic classification and statistical analysis

A web-based program called Usearch (version 7.0, http://drive5.com/uparse/) was applied to carry out taxon-dependent analysis. 16S rRNA gene sequences were used for phylogenetically consistent bacterial taxonomy according to the method of a Bayesian classifier, Ribosomal Database Project Classifier [41]. The Silva Database (Silva_128_16s, http://www.arb-silva.de) was compared to calculate the operational taxonomic units (OTUs) for all samples to show the abundance of bacterial species with 97% of identity cutoff, whereas the species for which the sum of OTUs of all the samples was more than 20 reads were retained. The richness of OTUs for each sample was produced at the level of genus. The length of all the valid bacterial sequences with primers was 440 bp on the average. The calculation of abundance at the level of genus was transformed according to log_2_ and normalized as the method of Niu et al. [42]. Inter-group OTUs were compared by the generation of a Venn diagram. The bacterial community indices adopted contained Chao and Shannon’s coverage. The diversity of bacteria was presented by the quantity of OTUs.

### *In vitro* rumen fermentation

The substrate was made with the feed of group M in the feeding experiment by drying and grinding through a 0.45 mm sieve. Collected from different dietary treatments during the feeding experiment, the ruminal fluid was filtrated by four-layer cheesecloth and mixed with pre-heated artificial saliva [43] at a ratio of 2:1 (buffer: ruminal fluid, v:v). The ruminal fluid (150 mL) which was buffered was dispensed into pre-warmed 200-mL incubation flasks. Two grams of each substrate was blended with the buffered ruminal fluid in each incubation flask which was incubated at the temperature of 39 °C for 12 h in water. The production ratio of methane was measured by real-time *in vitro* fermentation system (produced by Jilin Academy of Agricultural Sciences, code Qtfxy-6), which was tested for the effluent gas discharged from each incubation flask. The nitrogen (purity 99.99%) was passed into the incubation flask from the bottom at the speed of 200 mL/min. Methane was carried by nitrogen into an AGM10 sensor (Sensors Europe GmbH, Erkrath, FRG) and the concentration of methane was measured and recorded every 6 min [26]. The fermentation was terminated by placing flasks on ice. After opening the incubation flask, pH was measured using a PHS-3C pH meter (Shanghai INESA Scientific Instrument Co., Ltd., China), and 2 mL of incubation medium was collected for NH_3_-N [44]. Another 1 mL of incubation medium was analyzed for volatile fatty acids (VFAs), including acetic acid (AA), propionic acid (PA), and butyric acid (BA) using gas chromatography (Agilent Technologies 7890A GC System, USA) and the method of Castro-Montoya et al. [45]. The left fluid was dried in a forced-air oven at 60 °C for 72 h and placed in sealed containers in order to analyze the in vitro dry matter digestibility (IVDMD) [46].

### Experimental feeds and chemical analyses

Collected in plastic bags, the samples for diets were reserved at −20 °C. After the feeding experiment, the samples were warmed at 65 °C to a fixed weight. Thereafter, a 0.45 mm sieve and a high-speed universal pulverizer were used to grind them for analysis. The filter bag technique of ANKOM A200 (AOAC 973.18) was adopted to analyze neutral detergent fiber (NDF), acid detergent lignin (ADL) as well as acid detergent fiber (ADF). A Kjeltec 8400 analyzer unit (Foss, Sweden) was applied to measure the content of crude protein (CP) on the basis of the Kjeldahl method (AOAC 984.13). In addition, a Soxhlet apparatus was used to measure the content of ether extract (EE) based on Soxhlet extraction method (AOAC 920.85). Methods of Horwitz et al. [46] were the foundation of all chemical analyses.

### Data analysis

Sequences of good quality were deeply studied through its uploading to QIIME [39]. A comparison was made between valid bacterial sequences and sequences present in the Silva Database which classified the abundance calculation of each taxon with the optimal choice of classification [36]. QIIME filed the sequence length. Mothur was used for the generation of abundance and diversity indexes. After the implementation of a pseudo-single relevancy algorithm, there was 97% of OTUs identity cutoff [47–48]. For all the parameters, data was analyzed by the R-Studio software (version 7.2). Methane production was up to the approach of Sun et al. [26]. A one-way analysis of variance (ANOVA) was carried out late in each bioassay to compare selected taxonomic groups (abundant phyla or genera), bacterial community indices observed OTUs or methane production indices. Duncan’s test was adopted to perform the mean comparison at the significance level of P < 0.05. Redundancy analysis (RDA) was conducted to assess the association between the nutrients in the feed and the bacterial abundance in the rumen. The relationship among bacterial abundance, methane production, peak concentration (C_max_) and the time to peak concentration (T_max_) was assessed by means of Pearson’s correlations. All the data was presented as means ± S.E. (standard error).

## Results

### Relative quantification of total bacteria, methanogens, protozoa and anaerobic fungi

Firstly, the results of real-time qPCR (Table 3) showed that the numbers of methanogens and protozoa were increased with the decreasing F:C but not significantly. Conversely, the numbers of total bacteria and anaerobic fungi were decreased with the decreasing F:C. And the difference of total bacteria in three groups was significant, while the difference of anaerobic fungi was not. These results exhibited that the change of F:C has extremely effect only on the number of total bacteria but not on the other kinds of microorganism.

**Table 3.** Relative quantification of total bacteria, methanogens, protozoa and anaerobic fungi with different forage-to-concentrate ratio

| Item | Forage-to-concentrate ratio <sup>a</sup> |  |  | *P*-value |
| --- | --- | --- | --- | --- |
|  | L | M | H |  |
| Methanogens | 5.47±0.14 | 5.32±0.11 | 5.19±0.09 | 0.257 |
| Protozoa | 4.37±0.13 | 4.17±0.10 | 4.13±0.13 | 0.369 |
| Anaerobic fungi | 1.58±0.07 | 1.70±0.08 | 1.86±0.10 | 0.096 |
| General bacterial | 1.08±0.03 | 0.99±0.04 | 0.91±0.04 | 0.017 |

<sup>a</sup> L = forage-to-concentrate ratio 50:50; M = forage-to-concentrate ratio 70:30; H = forage-to-concentrate ratio 90:10. All the data are presented as mean ± S.E. (standard error).

### Analysis of DNA sequence data

After quality was controlled preliminarily, 517,492 paired-end 440-bp reads were obtained in total. Each sample got 28,750 sequences averagely. Reads had an overall length of 2.28 gigabases (GB), and each sample had a mean read length of 0.13 GB with 191,537, 171,125, and 154,794 raw reads in L, M and H groups respectively (Table 4). Based on 97% species similarity, 132,987, 104,640 and 92,194 OTUs were separately obtained from the samples in L, M and H groups (Table 4). Among all the samples, 708 OTUs were identified, of which 542 existing in all the groups were known as key OTUs (Fig 1A). Key OTUs took up about 76.6% of all OTUs, whereas 6, 5 and 11 OTUs were individually identified in groups L, M and H respectively. Good’s coverage was 99.4%, 99.3% as well as 99.2% for L, M and H groups separately, indicating the capture of dominant phylotypes by this study. The three groups were similar in diversity (Fig 1B). The richness (*P*<0.01) of the rumen microbiota was related to F:C (Fig 1C).

**Fig 1.**
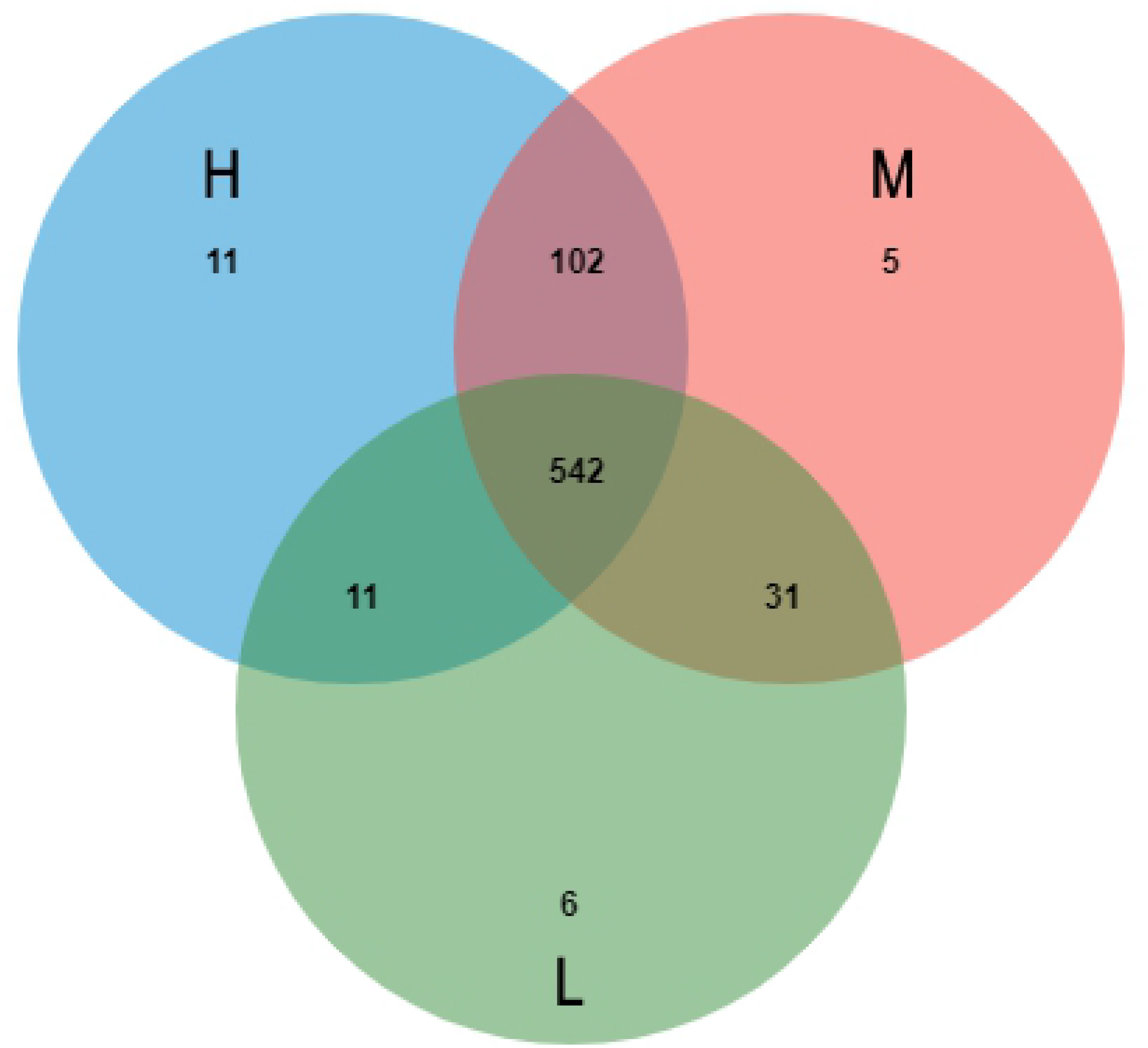
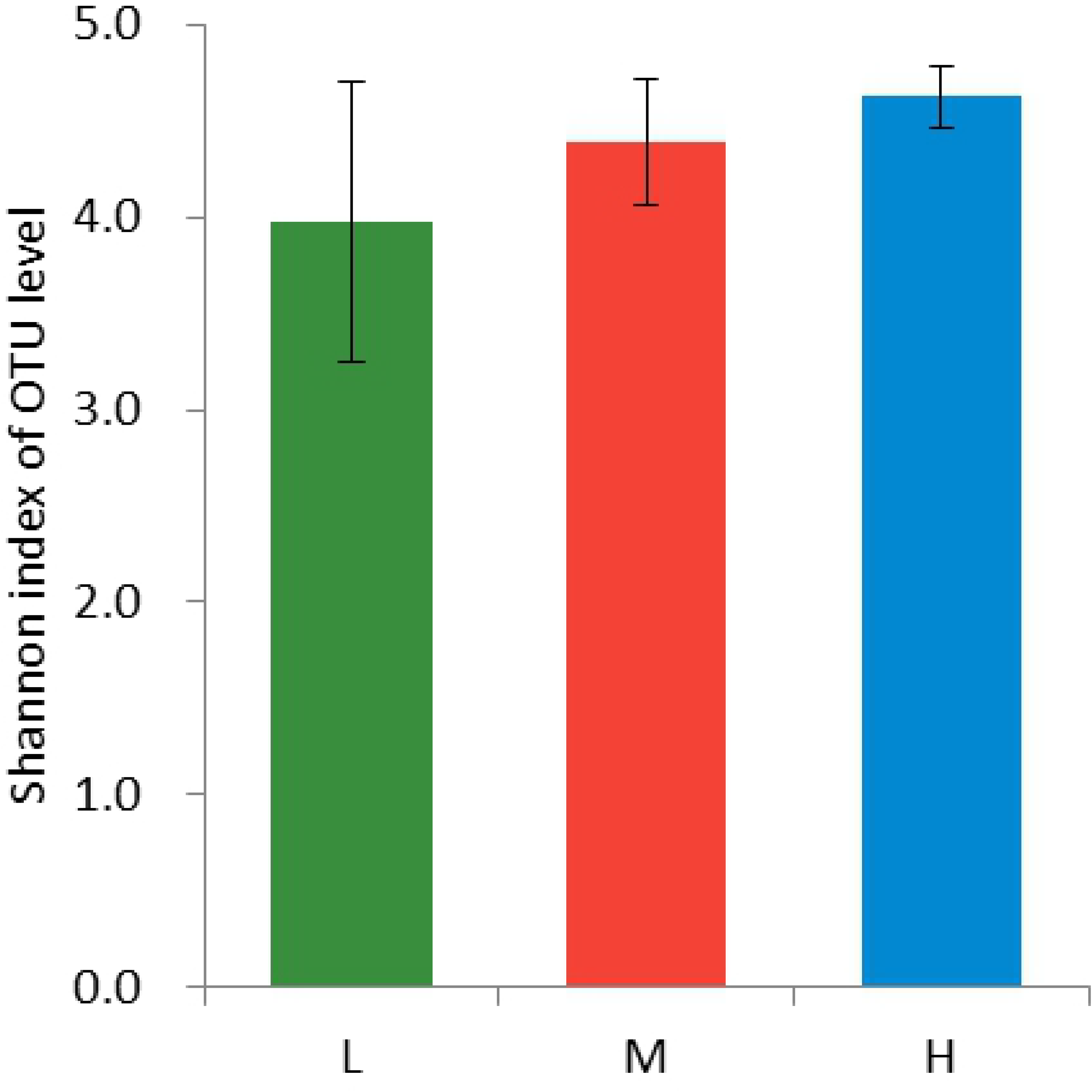
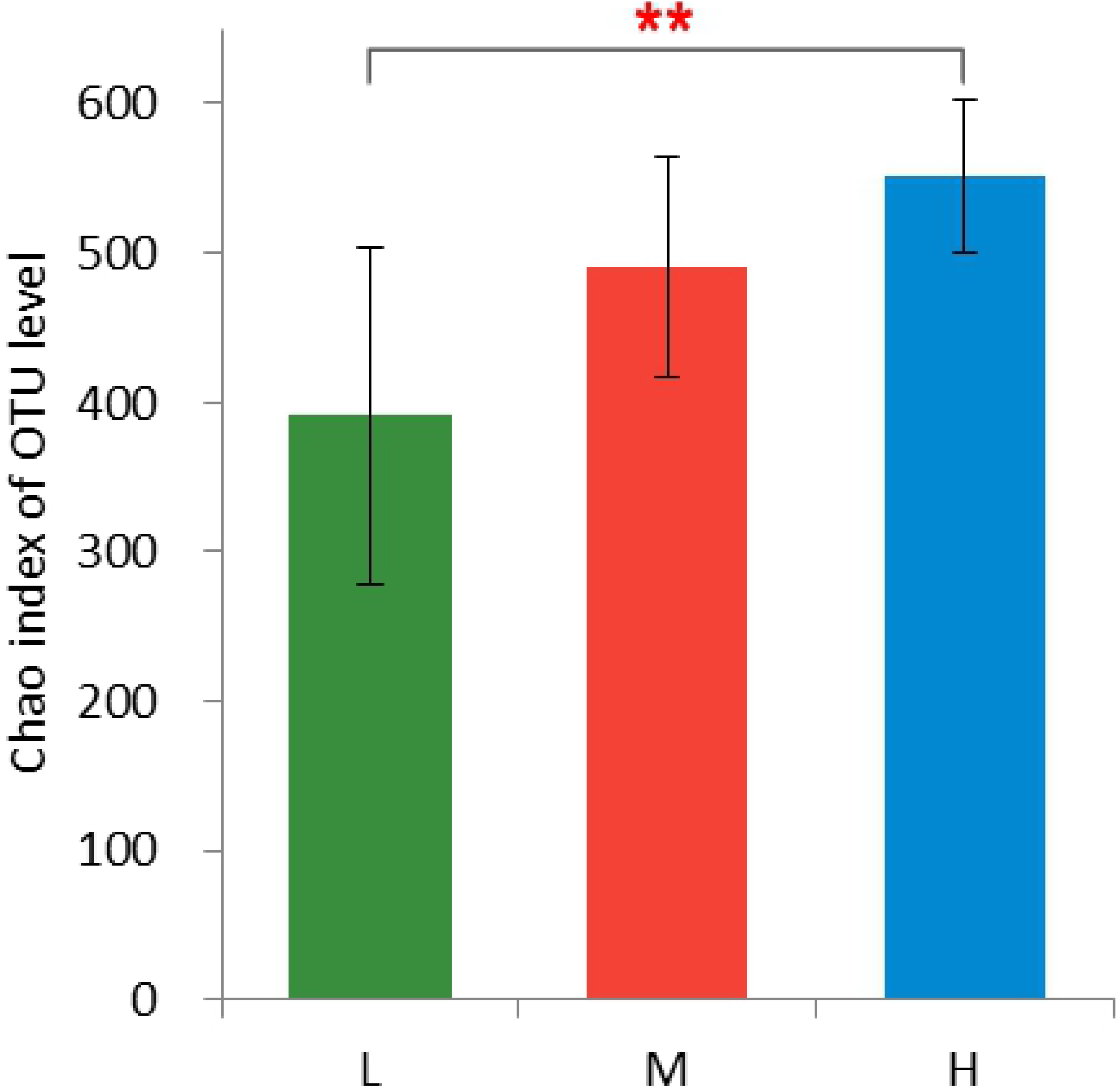
Comparison of the operational taxonomic units (OTUs) in the groups L, M, and H. OTUs, operational taxonomic units; L, forage-to-concentrate ratio, 50:50; M, forage-to-concentrate ratio, 70:30; H, forage-to-concentrate ratio, 90:10. The number of observed OTUs sharing ≥ 97% nucleotide sequence identity is shown (1A) Venn diagram showing the common and unique OTUs among the three groups. (1B) Bacterial diversity as determined from the Shannon index of OTUs in the three groups. (1C) Bacterial richness, as reflected in the Chao index.

**Table 4.** Raw reads and OTUs in the groups with different forage-to-concentrate ratio

| Group <sup>a</sup> | Raw reads | High quality reads | OTUs |
| --- | --- | --- | --- |
| L | 191,537 | 132,987 | 590 |
| M | 171,125 | 104,640 | 680 |
| H | 154,794 | 92,194 | 666 |
| Total | 517,492 | 329,821 | 1,936 |

OTUs, operational taxonomic units.
<sup>a</sup> L = forage-to-concentrate ratio 50:50; M = forage-to-concentrate ratio 70:30; H = forage-to-concentrate ratio 90:10.

### Bacterial community structure at the levels of phylum and genus

According to the results in Fig 2A, DNA sequences were distributed in different phyla. The three groups shared 14 phyla, namely *Bacteroidetes*, *Actinobacteria*, *Cyanobacteria*, *Chloroflexi*, *Elusimicrobia*, *Synergistetes*, *Fibrobacteres*, *Firmicutes*, *Lentisphaerae*, *Verrucomicrobia*, *Proteobacteria*, *Saccharibacteria*, *Spirochaetes* and *Tenericutes*. As the main components of the 14 phyla (*P*<0.01) in spite of the diet, *Bacteroidetes* and *Firmicutes* occupied over 90% of all sequences. The three groups showed differences in the bacterial richness of different phyla. The remarkable differences of bacterial richness in five out of 14 phyla were discovered in the three groups (Table 5). As the dominant phylum in group L, *Bacteroidetes* (*P*<0.01) accounted for about 66.14% of the sequences. Groups M and H assigned a lower percentage (60.05% and 56.80%) of the sequences to *Bacteroidetes*. Ranking the second as a phylum in all the groups, *Firmicutes* (*P*<0.01) comprised roughly of 24.13%, 27.45% and 32.01% sequences in the L, M and H groups respectively. The proportion of *Firmicutes* increased with the increase of the ratio. Moreover, the richness of *Proteobacteria*, *Spirochaetes* and *Synergistetes* changed with F:C (Table 5). With the increase of the ratio, the proportion of *Proteobacteria* and *Spirochaetes* (Table 5) decreased (*P*<0.01) and the proportion of *Synergistetes* (P < 0.01) (Table 5) increased.

**Fig 2.**
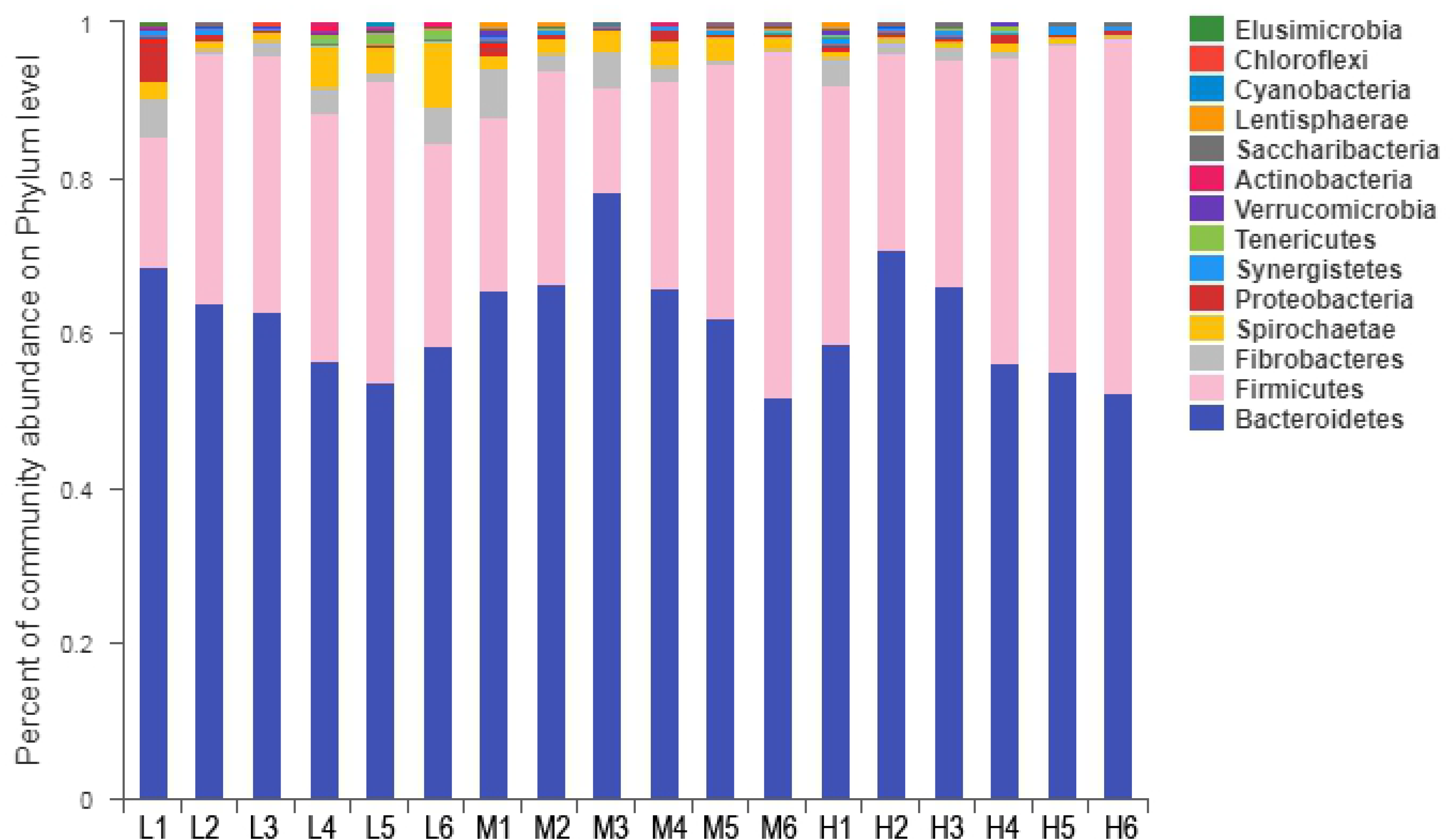
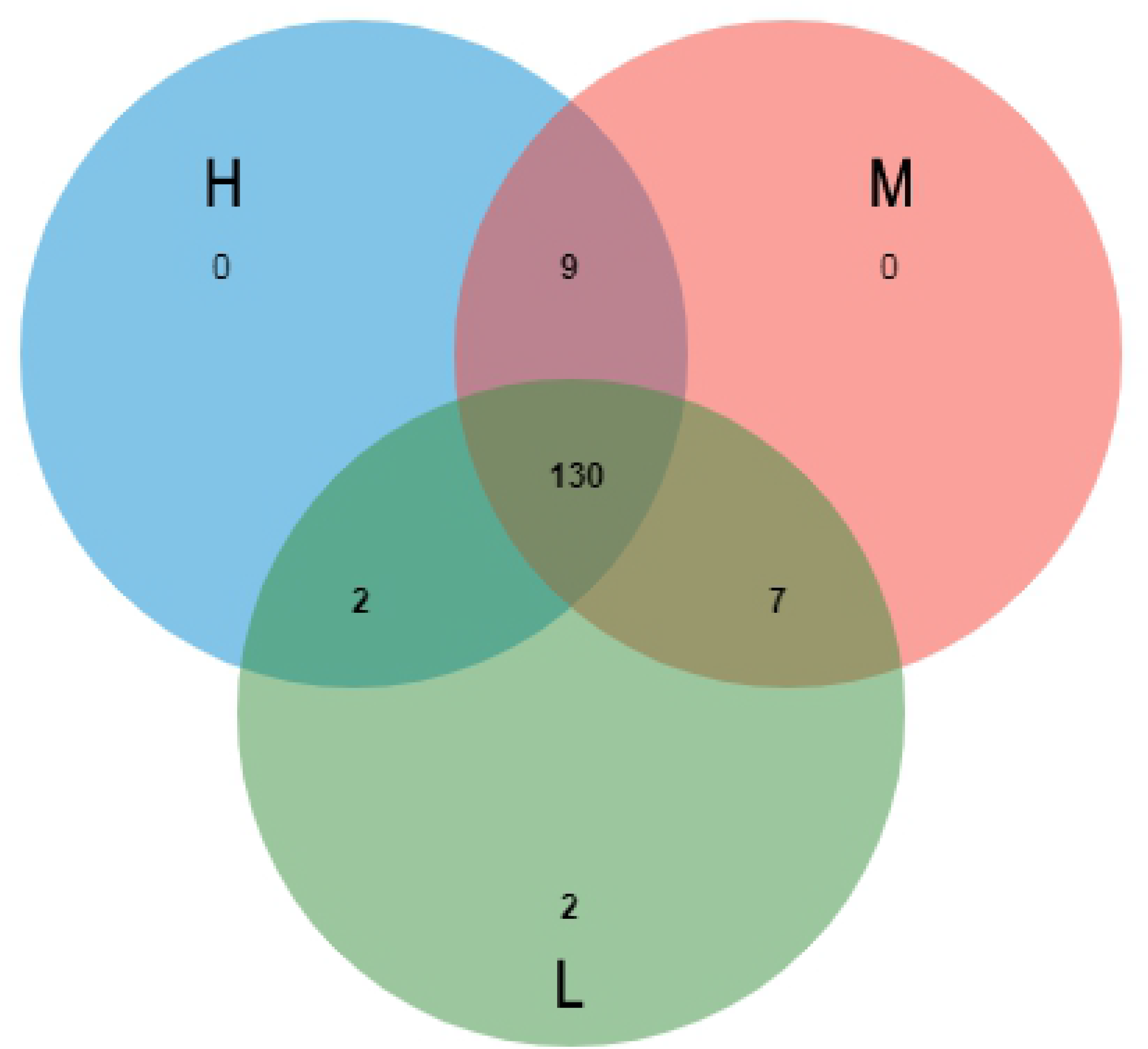
Phyla distribution of rumen flora and a Venn diagram of the genera in the groups of different forage-to-concentrate ratio. L, forage-to-concentrate ratio, 50:50; M, forage-to-concentrate ratio, 70:30; H, forage-to-concentrate ratio, 90:10. (2A) Distribution of the phyla as a percentage of the total number of identified 16S rDNA sequences in the groups of different forage-to-concentrate ratio. (2B) Venn diagram showing the comparison of genera between the groups at the same time points and depicting the genera that were unique to the three groups.

**Table 5.** Relative abundance of five distinct phyla and Pearson’s correlations in the groups with different forage-to-concentrate ratio

| Phylum | Forage-to-concentrate ratio <sup>a</sup> |  |  | Pearson's correlation <sup>b</sup> |
| --- | --- | --- | --- | --- |
|  | L | M | H |  |
| *Bacteroidetes* | 66.14±2.92 | 60.05±3.50 | 56.80±3.70 | -0.801** |
| *Firmicutes* | 24.13±2.87 | 27.45±2.92 | 32.01±2.56 | 0.814** |
| *Proteobacteria* | 1.70±0.13 | 0.77±0.07 | 0.59±0.03 | -0.920** |
| *Spirochaetae* | 3.02±0.26 | 2.41±0.26 | 0.75±0.05 | -0.951** |
| *Saccharibacteria* | 0.05±0.01 | 0.22±0.03 | 0.28±0.05 | 0.932** |

<sup>a</sup> L = forage-to-concentrate ratio 50:50; M = forage-to-concentrate ratio 70:30; H = forage-to-concentrate ratio 90:10. All the data are presented as mean ± S.E. (standard error).
<sup>b</sup> \* $P < 0.05$ , \*\* $P < 0.01$

At the level of genus, the identification of 150 genera in all the samples was conducted despite F:C (Fig 2A). L, M and H groups had 141, 146 and 141 genera respectively and shared 130 genera, whereas *Coriobacteriaceae* and *Gemella* were special for group L (Fig 2B). 15 richest genera, comprising over 78.96% of all sequences, included *Prevotella*, *Ruminococcus*, *Lachnospira*, *Rikenella*, *Succiniclasticum*, *Fibrobacter*, *Christensenella*, *Saccharofermentans*, *Eubacterium*, *Papillibacter*, *Quinella*, *Phocaeicola*, *Verllonella*, *Moryella* and *Fretibacterium*. The bacterial richness of 22 genera varied with the diet ratio. The abundance of 14 bacteria increased, whereas that of eight bacteria decreased (Table 6). Significantly different among all the groups, *Lachnospira*, *Fibrobacter* and *Clostridium* were not linearly related to the diet ratio. Among the linearly changed genera, *Prevotella* was the predominant genus, accounting for 50.79%, 43.06% and 34.09% of the total sequences in L, M and H groups respectively.

**Table 6.** Relative abundance of 25 distinct genera and Pearson’s correlations in the groups with different forage-to-concentrate ratio

| Genus | Forage-to-concentrate ratio <sup>a</sup> |  |  | Pearson's correlation <sup>b</sup> |
| --- | --- | --- | --- | --- |
|  | L | M | H |  |
| *Prevotella* | 50.79±2.53 | 43.06±4.48 | 34.09±4.65 | -0.901** |
| *Ruminococcus* | 5.03±0.49 | 9.18±1.34 | 10.33±0.76 | 0.902** |
| *Lachnospira* | 8.48±0.77 | 10.63±0.91 | 3.34±0.41 | -0.282 |
| *Rikenella* | 1.65±0.10 | 3.72±0.60 | 14.09±0.79 | 0.730* |
| *Succiniclasticum* | 4.11±0.50 | 5.49±0.29 | 7.04±0.39 | 0.963** |
| *Fibrobacter* | 2.53±0.05 | 2.89±0.11 | 1.52±0.10 | -0.603 |
| *Eubacterium* | 0.39±0.06 | 0.97±0.11 | 0.96±0.12 | 0.625 |
| *Papillibacter* | 0.05±0.01 | 0.46±0.05 | 1.41±0.15 | 0.768* |
| *Quinella* | 1.31±0.14 | 0.25±0.01 | 0.07±0.01 | -0.819* |
| *Verillonella* | 0.61±0.06 | 0.52±0.04 | 0.25±0.03 | -0.933** |
| *Fretibacterium* | 0.26±0.02 | 0.38±0.02 | 0.61±0.05 | 0.968** |
| *Anaerovorax* | 0.12±0.01 | 0.42±0.04 | 0.58±0.07 | 0.964** |
| *Pseudobutyrvibrio* | 0.04±0.01 | 0.27±0.02 | 0.77±0.03 | 0.976** |
| *Butyrvibrio* | 0.11±0.02 | 0.25±0.03 | 0.61±0.04 | 0.961** |
| *Ruminobacter* | 0.84±0.14 | 0.09±0.01 | 0.02±0.00 | -0.686* |
| *Selenomonas* | 0.13±0.01 | 0.15±0.02 | 0.61±0.09 | 0.734* |
| *Lachnoclostridium* | 0.18±0.01 | 0.20±0.02 | 0.50±0.03 | 0.788* |
| *Oribacterium* | 0.33±0.04 | 0.16±0.01 | 0.11±0.01 | -0.832* |
| *Syntrophococcus* | 0.38±0.05 | 0.11±0.01 | 0.01±0.00 | -0.957** |
| *Succinivibrio* | 0.22±0.03 | 0.18±0.01 | 0.13±0.01 | -0.893** |
| *Candidatus* | 0.05±0.00 | 0.19±0.02 | 0.24±0.04 | 0.926** |
| *Clostridium* | 0.17±0.01 | 0.25±0.02 | 0.04±0.00 | -0.631 |
| *Anaerotruncus* | 0.04±0.00 | 0.13±0.01 | 0.25±0.01 | 0.992** |
| *Olsenella* | 0.33±0.01 | 0.06±0.01 | 0.02±0.00 | -0.721* |
| *Ruminiclostridium* | 0.06±0.01 | 0.09±0.00 | 0.20±0.01 | 0.858** |

<sup>a</sup> L = forage-to-concentrate ratio 50:50; M = forage-to-concentrate ratio 70:30; H = forage-to-concentrate ratio 90:10. All the data are presented as mean ± S.E. (standard error).
<sup>b</sup> \* $P < 0.05$ , \*\* $P < 0.01$

### Nutrition index in rumen and its correlation with the rumen microbiota

In terms of the RDA, our dataset changed, which was principally interpreted by the increasing F:C (Fig 3). It suggested that 100% change in bacteria was explained by all the nutrition indices whose order of contribution was CP > ADF > NDF > Starch > EE > ADL (Table 7). The two sorting axes accounted for 95.48% of the changes based on this model with the first sorting axis explaining a change of 66.37% and 29.11% for the second sorting axis. The rumen microbiota in group L was concentrated in the regions with high CP, starch content and low NDF and ADF contents, whereas the rumen microbiota in group H was concentrated in the regions with high NDF and ADF contents and low CP and starch contents. The rumen microbiota in group M was concentrated in the regions with intermediate nutrient levels. According to the RDA analysis, the relevance of CP accounted for 0.72 of the microbiota (*P*<0.01) as the main nutrient factor affecting the structure of microbiota. Insignificant, the relevance of EE and ADL was the lowest (R^2^=0.41, *P*>0.05; R^2^=0.36, *P*>0.05) and they were not significant. Under the different levels of F:C, the different kinds of bacteria bacterial community were established, which could reflect the state of the microflora in the rumen fluid at the beginning of *in vitro* fermentation.

**Fig 3.**
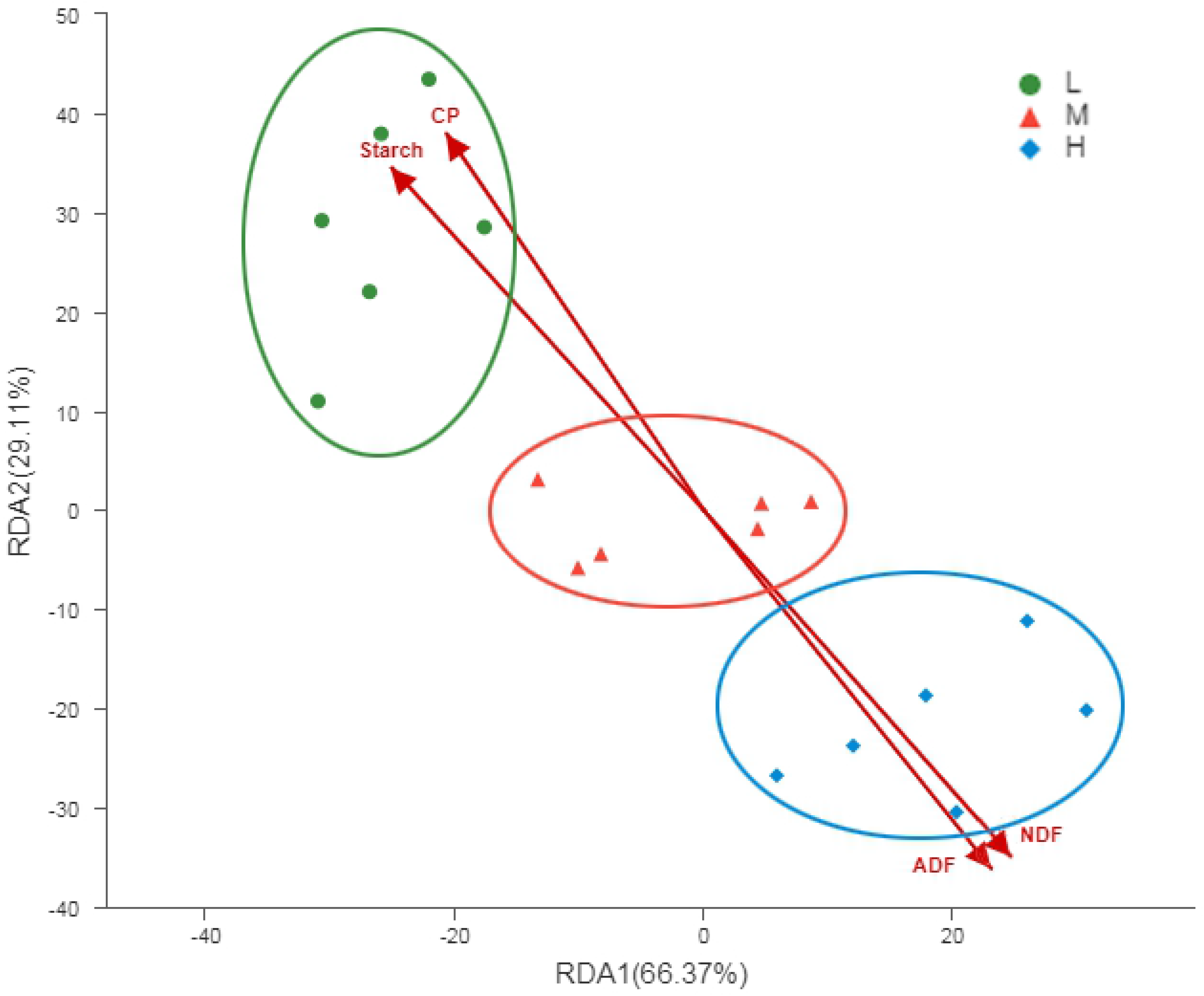
Redundancy analysis of nutrition index and the rumen microbiota in the groups of different forage-to-concentrate ratio. L, forage-to-concentrate ratio, 50:50; M, forage-to-concentrate ratio, 70:30; H, forage-to-concentrate ratio, 90:10. Two sorting axes accounted for 95.48% of the changes with the first sorting axis explaining a change of 66.37% and 29.11% for the second sorting axis.

**Table 7.** Relevance of nutrition indices in the redundancy analysis

| Item | RDA<sub>1</sub> | RDA<sub>2</sub> | R<sup>2</sup> |
| --- | --- | --- | --- |
| CP | -0.4325 | 0.9016 | 0.72** |
| ADF | 0.4867 | -0.8736 | 0.70** |
| NDF | 0.5225 | -0.8527 | 0.69** |
| Starch | -0.5321 | 0.8467 | 0.69** |
| EE | 0.4641 | 0.3559 | 0.41 |
| ADL | 0.3482 | -0.372 | 0.36 |

CP = crude protein, ADF = acid detergent fiber, NDF = neutral detergent fiber, EE = ether
extract, ADL = acid detergent lignin.
\* $P < 0.05$ , \*\* $P < 0.01$

### *In vitro* rumen fermentation characteristics, real-time methane production and its correlation with the rumen microbiota

After 12 h fermentation, the concentrations of pH, AA and A/P were decreased greatly with the decreasing F:C. Simultaneously, the concentrations of PA, BA, NH_3_-N, and IVDMD were increased with the decreasing F:C. The greatly growth of IVDMD had led to the massive production of VFAs (Table 8). The C_max_ (*P*<0.01, Table 9, Fig 4) and total production (*P*<0.05, Table 9) of methane decreased with the increase in F:C, whereas T_max_ (*P*<0.01, Table 9, Fig 4) of methane increased with the increase of F:C. At the level of phylum, *Bacteroidetes*, *Proteobacteria* as well as *Spirochaetae* showed a positive correlation with the C_max_ and total production, and a negative correlation with T_max_ (Table 10). *Firmicutes* and *Saccharibacteria* were positively correlated with T_max_, but negatively correlated with C_max_ and total production (Table 10). At the level of genus, *Prevotella*, *Quinella*, *Verllonella*, *Ruminobacter*, *Oribacterium*, *Succinivibrio*, *Syntrophococcus* and *Olsenella* was positively correlated with C_max_ and total production in bacterial abundance, but negatively correlated with T_max_ (Table 11). *Ruminococcus*, *Rikenella*, *Succiniclasticum*, *Eubacterium*, *Papillibacter*, *Pseudobutyrivibrio*, *Butyrivibrio*, *Candidatus*, *Anaerotruncus* and *Ruminiclostridium* was positively correlated with T_max_ in bacterial abundance, but negatively correlated with C_max_ and total production (Table 11).

**Fig. 4.**
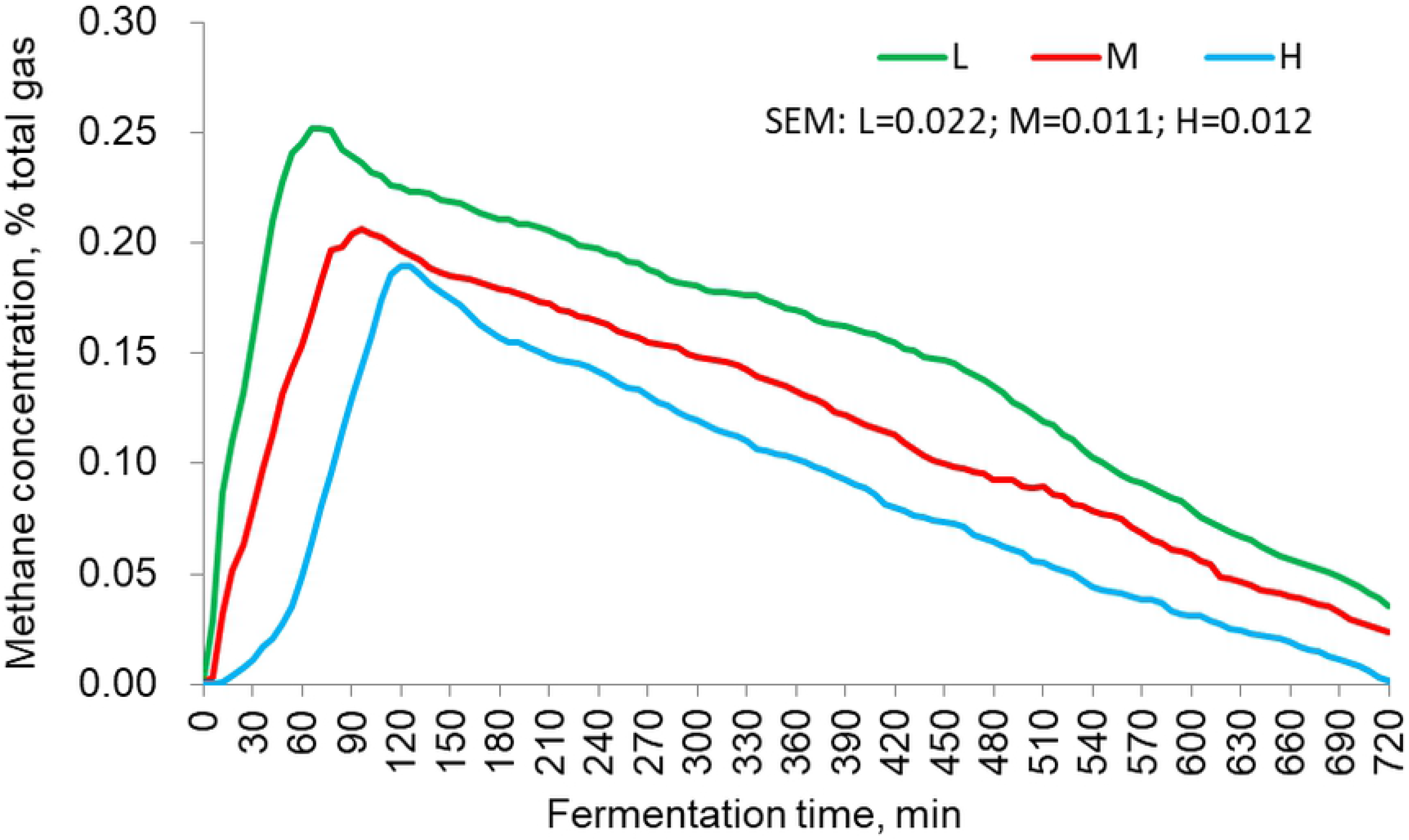
Methane production curve *in vitro* with different forage-to-concentrate ratio in the diets. L, forage-to-concentrate ratio, 50:50; M, forage-to-concentrate ratio, 70:30; H, forage-to-concentrate ratio, 90:10. SEM = standard error of mean.

**Table 8.** Effect of different forage-to-concentrate ratio on *in vitro* fermentation

| Item | Forage-to-concentrate ratio <sup>a</sup> |  |  | P-value |
| --- | --- | --- | --- | --- |
|  | L | M | H |  |
| pH | 6.66±0.09 | 6.84±0.10 | 6.95±0.11 | 0.048 |
| NH<sub>3</sub>-N | 29.51±0.75 | 28.97±0.71 | 26.80±0.59 | 0.025 |
| Acetate acid | 47.98±0.90 | 49.54±0.66 | 51.71±1.17 | 0.031 |
| Propionate acid | 15.70±0.96 | 13.75±0.67 | 12.11±0.80 | 0.018 |
| Butyrate acid | 8.64±0.18 | 8.04±0.16 | 7.95±0.26 | 0.049 |
| A/P | 2.93±0.14 | 3.40±0.13 | 4.02±0.16 | 0.004 |
| IVDMD | 68.04±1.91 | 62.16±1.14 | 60.51±2.15 | 0.016 |

A/P, Acetate acid/ Propionate acid; IVDMD, *in vitro* dry matter digestibility.
<sup>a</sup> L = forage-to-concentrate ratio 50:50; M = forage-to-concentrate ratio 70:30; H = forage-to-concentrate ratio 9:1. All the data are presented as mean ± S.E. (standard error).

**Table 9.** C_max_, T_max_ and total production of methane *in vitro* and Pearson’s correlations in the groups with different forage-to-concentrate ratio

| Item | Forage-to-concentrate ratio <sup>a</sup> |  |  | Pearson's correlation <sup>b</sup> |
| --- | --- | --- | --- | --- |
|  | L | M | H |  |
| C<sub>max</sub>, % | 0.25±0.01 | 0.21±0.02 | 0.19±0.01 | -0.827** |
| T<sub>max</sub>, min | 70.50±5.74 | 96.00±4.90 | 123.00±13.46 | 0.922** |
| Total production, mmol/g | 35.16±2.34 | 26.51±0.99 | 18.6±2.00 | -0.772* |

C<sub>max</sub>, peak concentration; T<sub>max</sub>, the time to peak concentration.
<sup>a</sup> L = forage-to-concentrate ratio 50:50; M = forage-to-concentrate ratio 70:30; H = forage-to-concentrate ratio 90:10. All the data are presented as mean ± S.E. (standard error).
<sup>b</sup> \**P* < 0.05, \*\**P* < 0.01

**Table 10.** Pearson’s correlations between five distinct phyla and *in vitro* C_max_, T_max_ and total production of methane with different forage-to-concentrate ratio

| Phylum | Pearson's correlations <sup>a</sup> |  |  |
| --- | --- | --- | --- |
|  | C<sub>max</sub> | T<sub>max</sub> | Total production |
| *Bacteroidetes* | 0.627* | -0.560* | 0.503 |
| *Firmicutes* | -0.614* | 0.579* | -0.500 |
| *Proteobacteria* | 0.563* | -0.727** | 0.587* |
| *Spirochaetae* | 0.620* | -0.732** | 0.738** |
| *Saccharibacteria* | -0.594* | 0.723** | -0.737** |

C<sub>max</sub>, peak concentration; T<sub>max</sub>, the time to peak concentration.
<sup>a</sup> \**P* < 0.05, \*\**P* < 0.01

**Table 11.**
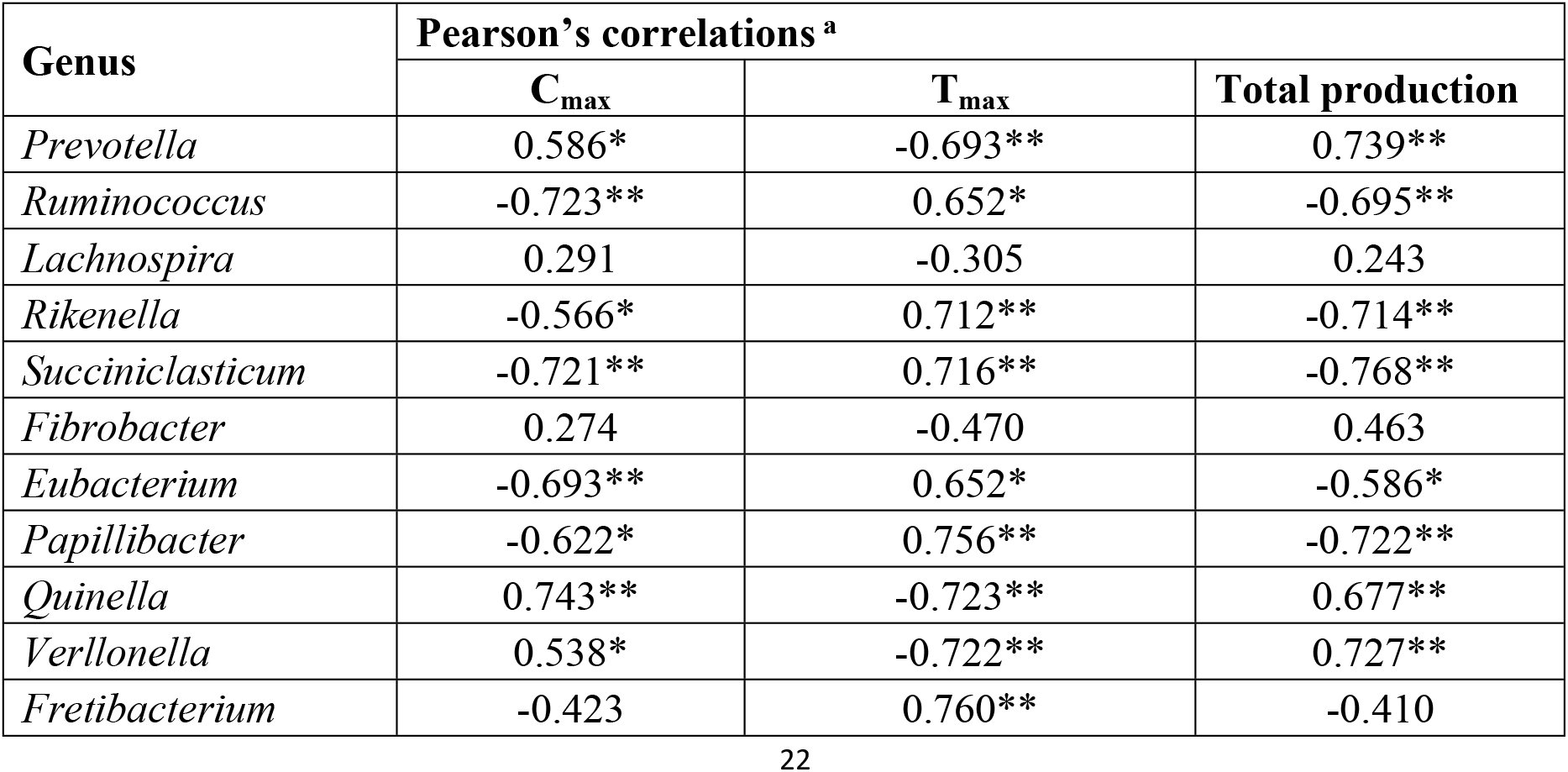
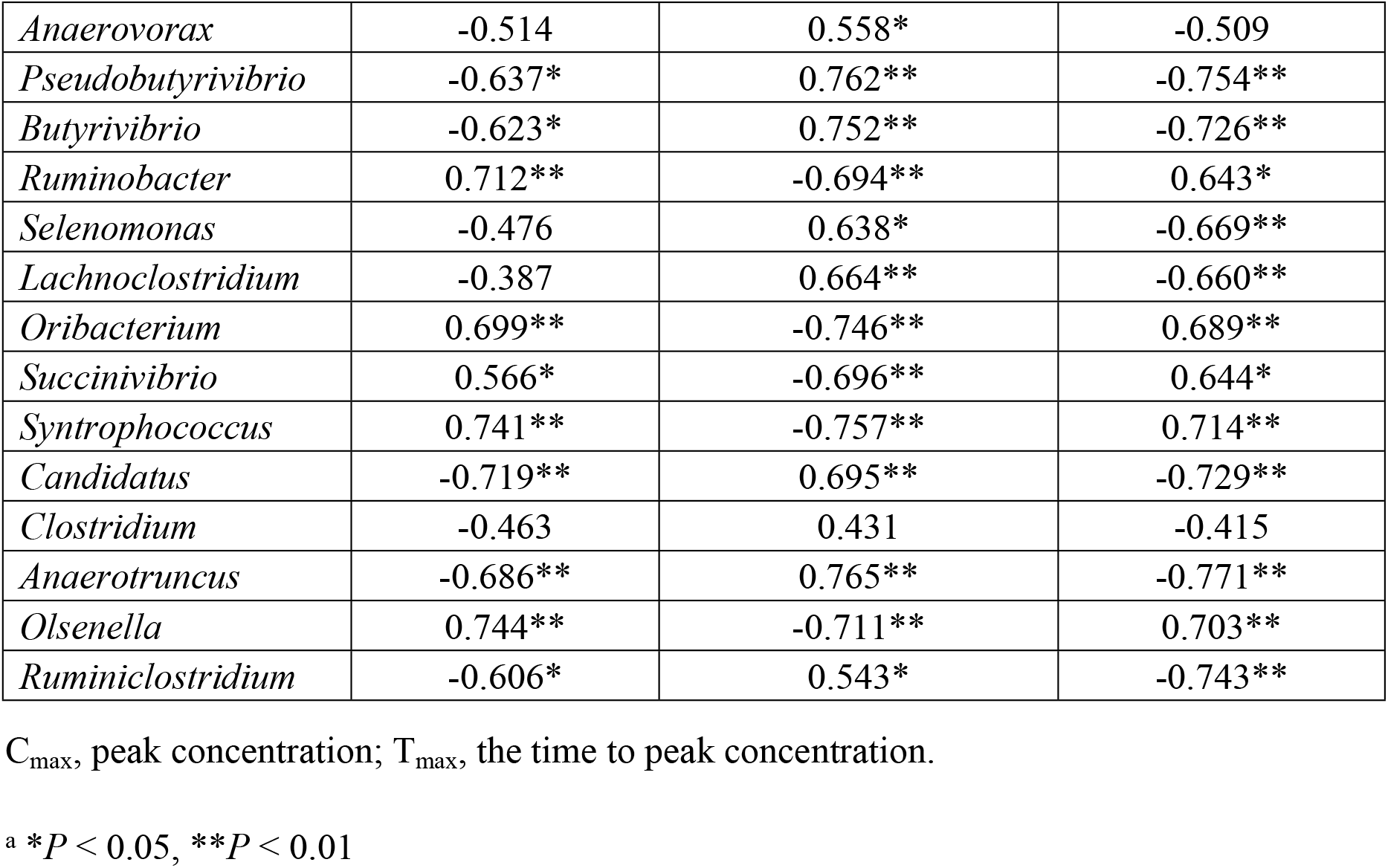
Pearson’s correlations between 25 distinct genera and *in vitro* C_max_, T_max_ and total production of methane with different forage-to-concentrate ratio

## Discussion

The C_max_ (*P*<0.01, Table 9, Fig 4) and total production (*P*<0.05, Table 9) of methane decreased with the increase in F:C, whereas T_max_ (*P*<0.01, Table 9, Fig 4) of methane increased with the increase of F:C. At the level of phylum, *Bacteroidetes*, *Proteobacteria* as well as *Spirochaetae* showed a positive correlation with the C_max_ and total production, and a negative correlation with T_max_ (Table 10). *Firmicutes* and *Saccharibacteria* were positively correlated with T_max_, but negatively correlated with C_max_ and total production (Table 10). At the level of genus, *Prevotella*, *Quinella*, *Verllonella*, *Ruminobacter*, *Oribacterium*, *Succinivibrio*, *Syntrophococcus* and *Olsenella* was positively correlated with C_max_ and total production in bacterial abundance, but negatively correlated with T_max_ (Table 11). *Ruminococcus*, *Rikenella*, *Succiniclasticum*, *Eubacterium*, *Papillibacter*, *Pseudobutyrivibrio*, *Butyrivibrio*, *Candidatus*, *Anaerotruncus* and *Ruminiclostridium* was positively correlated with T_max_ in bacterial abundance, but negatively correlated with C_max_ and total production (Table 11).

Firstly, the results of real-time qPCR showed that the number of methanogens, protozoa and anaerobic fungi, under the change of F:C, was changed significantly except bacterial. Archaea was considered the producer of methane. But with the change of F:C in this study, the quantity of archaea was stable. It showed no significant linear relationship between the structure of archaea and methane production, which is similar to the research of Lengowski et al. [25]. The relationship between archaea and methane production has been discussed in many studies. Moreover, the main raw materials for methane synthesis, such as hydrogen, carbon dioxide and volatile fatty acids, were produced by bacteria [49]. Methane synthesis was a passive behavior of archaea to maintain rumen pressure and pH balance in the case of too high ratios of bacterial synthesis [50–51]. Therefore, methane production was more probably related to the concentration of synthetic raw materials in the rumen and the bacteria producing these materials.

The second goal of this experiment was to explore changes in the genus level of the rumen flora with different F:C. With the increase of the ratio, the proportion of different genera showed significant differences, revealing the effectiveness of experimental gradient design. In this study, the proportion of *Prevotella* showed a linearly increasing trend with the increase of protein levels in diets, which was consistent with the results of Xu and Gordon [52]. As a genus, *Prevotella* has many functions, mainly including promoting protein degradation and assisting other strains in enhancing the utilization of fiber materials in ruminants [53]. *Ruminococcus*, a cellulolytic bacterium [54], increased with the increasing fiber. *Succinivibrio*, *Ruminobacter*, *amylophilus* and *Selenomonas* were starch-degrading bacteria that could produce acetic acid and succinic acid during starch degradation [55]. Succinic acid was eventually transformed to PA [56] to provide energy for microbial protein synthesis in the rumen. With the decrease of starch content in the diets, the proportion of these three genera decreased significantly in this experiment, indicating the change of carbohydrate fermentation substrate from a non-structural carbohydrate to a structural carbohydrate. *Butyrivibrio* and *Pseudobutyrivibrio* were carbohydrate-degrading bacteria producing butyric acid [57]. In this experiment, the proportion of *Butyrivibrio* and *Pseudobutyrivibrio* decreased linearly with the increase of starch, but increased linearly with the increase of NDF and ADF in the diets, which showed that *Butyrivibrio* and *Pseudobutyrivibrio* were more likely to produce energy by using structural carbohydrates. The proportion of *Eubacterium* with the function of degrading structural carbohydrates was similar to that of *Butyrivibrio* [58].

In this study, the third aim was to gain a preliminary understanding of the relationship between nutrition levels and the diversity and richness of rumen microbiota in sheep under various F:C. In this experiment, CP was the most important nutrient factor contributing to the change in bacterial diversity. Bodine and Purvis [59] found that the effect of supplementation of non-structural carbohydrate is largely determined by the level of protein in the diet. Adding protein to the diet can improve the balance of energy and nitrogen and increase digestibility. CP could provide nitrogen resource for the self-replication and enzyme synthesis of bacteria [60]. ADF, NDF and starch were important nutrient factors providing carbon resource for self-replication and energy. However, starch showed a negative correlation with bacterial diversity compared to ADF and NDF because of its easier decomposition as a non-structural carbohydrate. According to the studies of Kononoff and Heinrichs [61] and Drackley et al. [62], the rumen fermentation was mainly in the AA-mode when NDF and ADF contents in the diet were high and mainly in the PA-mode when starch content in the diet was high. The Chao index of OTUs increased with the increase in F:C, showing more strains of bacteria were required by the degradation of NDF and ADF to cooperate than those required by the degradation of CP and starch. These were the changes of microbiota in the rumen. EE and ADL in the diets had no significant effects on the changes in the rumen microbiota. Jenkins [63] found that only about 8% of fat in the rumen was degraded. There might be two reasons: The designed levels of EE and ADL content in the diets were too close to result in the similarity of the microbial community or these nutrients were not main energy resources for bacterial activity in the rumen so that bacteria were not sensitive to the low levels of EE [64]. Based on the above results, three kinds of rumen bacterial community were proved to be successfully established.

In this study, the final goal was to preliminarily understand the relationship between methane production and the rich and diversity of rumen microbiota in sheep under various F:C. In the anaerobic environment of the rumen, a variety of organic compounds could eventually be transformed to methane through decomposed by a number of microorganisms [5]. Leng and Nolan [14] showed that 80% of the nitrogen available to ruminal bacteria came from ammonia and 20% from amino acids or oligopeptides. With the increase of F:C, C_max_ of methane was delayed. For the diet with higher CP and starch contents, methane production could reach C_max_ more quickly, showing a significant correlation with the rumen microbiota. With lower CP content in diet, bacteria required more time for protein decomposition to provide materials for their reproduction and methane synthesis, which indicated that methane synthesis needed to go through a “start-up” phase before the normal fermentation mode. Previous studies only found that C_max_ of methane occurred at about 2 h after food intake [65–66]. The missing of the delay phenomenon could be attributed to the insufficient frequency of detection. On the other hand, both C_max_ and total production of methane with lower CP and starch contents in the diet during fermentation were less than those in the diets with lower NDF and ADF contents. It was probably because related bacteria like *Prevotella* and *Butyrivibrio* could decompose nitrogen compounds to provide sufficient raw materials for the reproduction and synthesis of methanogenic archaea [67]. As indicated by the correlation analysis, fiber-degrading bacteria were positively correlated with T_max_ of methane, but negatively correlated with C_max_ and total production of methane. Compared with fiber-degrading bacteria, starch-degrading and protein-degrading bacteria showed an opposite correlation. More readily available nitrogen and degradable carbohydrates could be preferentially used by microorganisms [68], providing more effective support for methane synthesis. Furthermore, C_max_ and T_max_ of methane could be effective parameters for predicting the type of rumen fermentation, which however remained to be confirmed by further research.

Based on nutritional parameters, a new model of methane prediction with a wider range of applications is being developed in accordance with the results of the present study. The genera of bacterial, as the parameters for the prediction models, had been narrowed down. There were significant correlations between specific bacterial at the starting of fermentation and real-time methane production *in vitro*. However, the dynamic changes of bacterial at the moment such as T_max_ during the fermentation need to be explored in the following study. Additionally, the fermentation in vivo was more complex. For instance, nitrogen cycling in ruminants might provide bacteria with initial nitrogenous material [69]. Thus, further studies are required to confirm the occurrence of this delay phenomenon in vivo and illustrate the process.

## Conclusions

With the change in F:C, bacterial community structure and methane production in the rumen showed significant changes. Crude protein was the most important nutrient factor that contributed to the change in bacterial diversity. Among the 150 genera identified in the rumen, the abundance of 22 varied linearly with F:C. These genera would be further screened to serve as effective parameters for the methane prediction model. In addition, during the 12 h *in vitro* fermentation, as F:C increased, the C_max_ and total production of methane decreased significantly, and T_max_ was delayed by 26–27 min. The fiber-degrading bacteria were positively correlated with this phenomenon, but starch-degrading and protein-degrading bacteria were negatively correlated with it.

## Acknowledgments

We are grateful to the technical staff of the Animal Nutrition and Feed Science Laboratory at Jilin Agricultural University (Changchun, China) for their help in this work. The *in vitro* fermentation system was provided by Animal Nutrition Institute of Jilin Academy of Agricultural Sciences (Changchun, China). The work on high-throughput sequencing was outsourced to Shanghai Majorbio Technology Co., Ltd.

